# Real-time visualization and quantification of human Cytomegalovirus replication in living cells using the ANCHOR DNA labeling technology

**DOI:** 10.1101/300111

**Authors:** Bernard Mariamé, Sandrine Kappler-Gratias, Martin Kappler, Stéphanie Balor, Franck Gallardo, Kerstin Bystricky

## Abstract

Human cytomegalovirus (HCMV) induces latent life-long infections in all human populations. Depending on geographic area and socio-economic conditions between 30 to nearly 100% of individuals are affected. The biology of this virus is difficult to explore due to its extreme sophistication and the lack of pertinent animal model. Here we present the first application of the ANCHOR DNA labeling system to a herpes virus, allowing real time imaging and direct monitoring of HCMV infection and replication in human living cells. The ANCHOR system is composed of a protein (OR) which specifically binds to a short, non-repetitive DNA target sequence (ANCH) and spreads onto neighboring sequences due to protein oligomerization. If OR protein is fused to GFP, this accumulation results in a site specific fluorescent focus. We have created a recombinant ANCHOR-HCMV harboring an ANCH target sequence and the gene encoding the cognate OR-GFP fusion protein. Infection of permissive cells with ANCHOR-HCMV enables visualization of the nearly complete viral cycle until cell fragmentation and death. Quantitative analysis of infection kinetics and of viral DNA replication revealed cell-type specific behavior of HCMV and sensitivity to inhibitors. Our results show that the ANCHOR technology is a very efficient tool for the study of complex DNA viruses and new highly promising biotechnology applications.

**IMPORTANCE:** The ANCHOR technology is to date the most powerful tool to follow and quantify the replication of HCMV in living cells and to gain new insights into its biology. This technology is applicable to virtually any DNA virus or virus presenting a dsDNA phase, paving the way to infection imaging in various cell lines or even in animal models and opening fascinating fundamental and applied prospects. Associated to high content automated microscopy, this technology permitted rapid, robust and precise determination of Ganciclovir IC50 and IC90 on HCMV replication, with minimal hands-on investment. To search for new antiviral activities, the experiment is easy to up-grade towards efficient and cost-effective screening of large chemical libraries. The simple infection of permissive cells with ANCHOR-viruses in the presence of a compound of interest may even provide a first estimation about the stage of the viral cycle this molecule is acting upon.

## INTRODUCTION

Human cytomegalovirus (HCMV), also called Human Herpesvirus 5 (HHV5), belongs to the β-*herpesviridae*family and, as all herpesviruses (HV), is able to establish life-long latency in infected individuals (1). HCMV is the largest HHV with a double stranded DNA genome of about 240kb. It is usually transmitted through body fluids such as saliva, urine or breast milk but also through sexual contacts (2). Primary infection is generally benign or silent in healthy individuals but may be much more serious and even life threatening in immuno-compromised patients, especially those having received hematopoietic cells or solid organ transplants, or AIDS patients. The virus is also able to cross the placental barrier and primary HCMV infection during pregnancy, especially during the first quarter, is the leading cause of birth defects, with an estimation of one million HCMV congenital infections worldwide per year (3, 4). Among these infections, possibly up to 25% of newborns will keep sensorineural and intellectual deficits. *In vivo* infection is poorly understood but most likely initiates in a mucosal tissue and then spreads through blood monocytes which disseminate the virus in numerous susceptible sites. HCMV binds to heparan sulfate proteoglycan (5) and to numerous cell membrane structures among which CD13 (6), annexin II (7), DC-SIGN (8), EGFR (9) and PDGFR-α (10) are candidate receptors. This may in part explain the remarkably broad cell tropism of this virus which is able to infect and replicate in numerous cell types including epithelial, dendritic, fibroblastic, endothelial or smooth muscle cells (11) and to establish latency in CD34+ hematopoietic progenitor cells (12). Long lasting efforts have allowed partial deciphering of the biology of this highly sophisticated virus but much remains to be done with regard to *in vivo* infection kinetics. Techniques to track real time infections in live cells have been developed for RNA viruses (13, 14, 15) and also for Herpes viruses (16, 17, 18). However, up to now, fluorescent tracking of HVs relied on simple GFP expression or on fusion of the GFP gene with a structural viral gene. These engineered viruses have greatly contributed to some pioneering work but did not provide quantitative information about replication kinetics of the viral genome. Therefore, to gain a better understanding of the fundamental biology of HVs, we have introduced a new technology enabling real time follow-up and counting of viral genomes during infection in live cells and also possibly in live animal models. In this paper, we present the use of the patented ANCHOR DNA labeling technology (19) for tracking of HCMV in living cells. ANCHOR is a bipartite system derived from a bacterial parABS chromosome segregation machinery. Under its natural form in bacteria, the parABS system consists in a short non repetitive target DNA sequence containing a limited number of nucleation parS sites to which the parB proteins bind and then spread onto adjacent DNA through a mechanism of protein-protein interaction. The third component of the system is an ATPase involved in the last steps of bacterial chromosomes or plasmids segregation. Under its engineered form, called ANCHOR, the OR protein (ParB) specifically binds to the cognate, shortened, ANCH sequence, which comprises palindromic parS nucleation sites (20, 21). If the OR protein is fused to a fluorescent protein (FP), its accumulation on the ANCH target sequence and spreading over neighboring sequences result in the formation of an easily detectable fluorescent focus, thereby identifying the position of the ANCH tagged DNA locus (Fig.1a). Different ANCHOR systems (1 to 4, derived from various bacteria) have been used successfully to analyze motion of single genomic loci and DNA double strand break processing in living yeast (22) and chromatin dynamics during transcription in human cells (23). These ANCHOR systems were shown not to perturb chromatin structure and function despite the presence of up to 500 OR proteins on and around the ANCH sequence (23). Here, we have created HCMV genomes containing the ANCH2 target sequence (HCMV-ANCH2) or both the ANCH3 target sequence and the gene encoding the corresponding OR3-GFP protein (HCMV-ANCHOR3). In the latter case, OR3-GFP proteins (which do not present any known intracellular localization sequence) freely diffuse in the cell and rapidly associate with the ANCH3 sequence, rendering the HCMV DNA fluorescent and detectable by microscopy as well defined spots over a uniform background of OR-GFP proteins. Thanks to these engineered virions, we were able to visualize early infection and initial duplication of the incoming genomes, viral DNA amplification, replication and cell death in real time and in live cells. All these steps were simply observed by microscopic examination, with no additional manipulation and without fixation, extraction or reagents of any kind, emphasizing the ease in use, the power and potential of this technology. Furthermore, analyzing the effect of Ganciclovir on ANCHOR-HCMV infection illustrates the remarkable potential of this technology for time and cost-effective screening of compound libraries in the search of new antiviral molecules. Its suitability for labeling any DNA virus (and possibly any virus presenting a dsDNA phase) offers unprecedented opportunities for new biotechnology applications.

**Figure 1.**
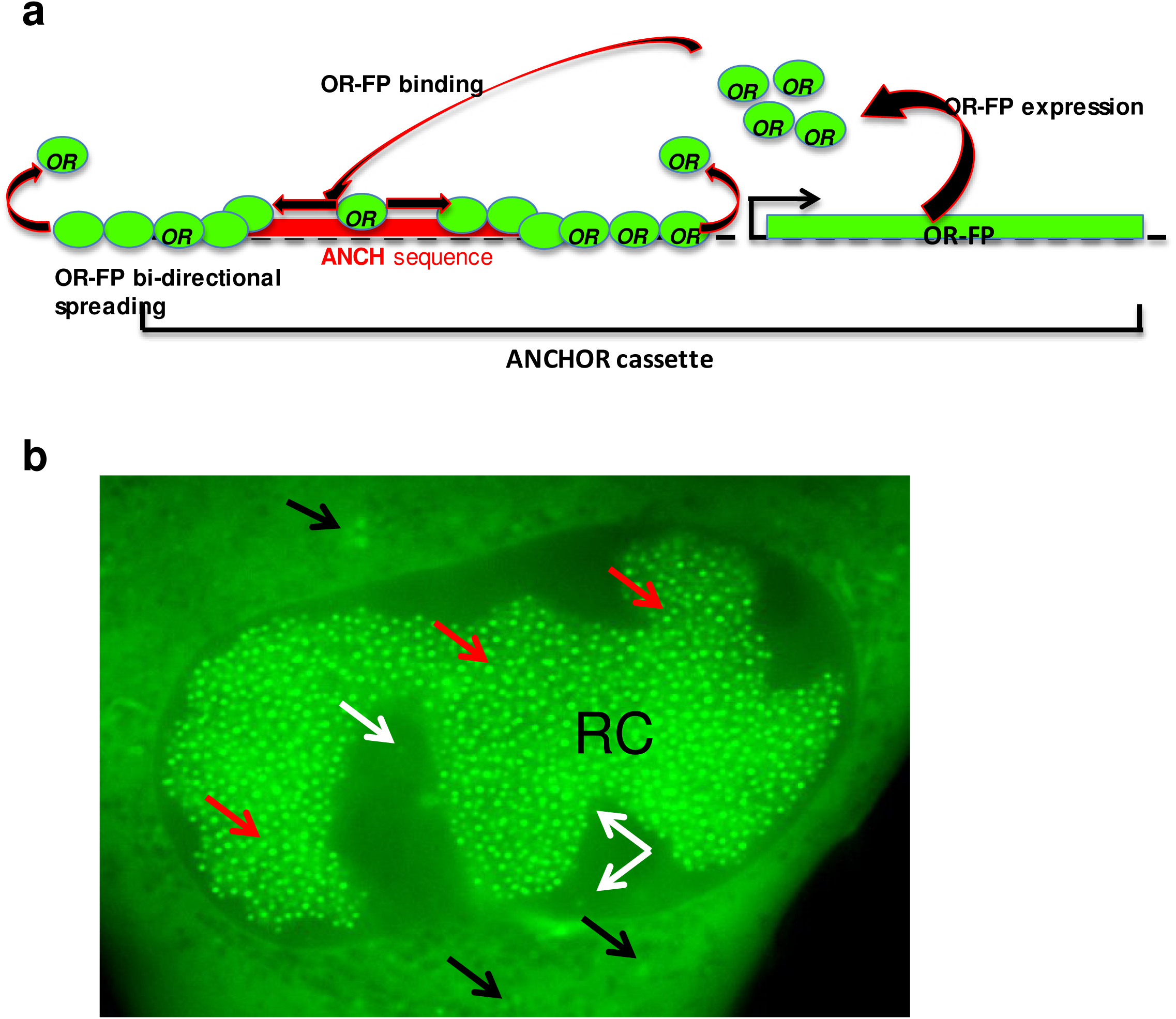
Principle of the ANCHOR DNA labeling technology. the ANCHOR system is composed of an ANCH DNA target sequence less than 1kb long (red box) which specifically binds dimers of OR protein through nucleation sites. Bound dimers spread on the DNA while further OR dimers are recruited to form a large metastable nucleoprotein complex. When OR protein is fused to a fluorescent marker (green circles), accumulation of this complex on the ANCH sequence forms a spot which is easily detected by fluorescence microscopy; b) TB40-ANCHOR3 infection of MRC5 cells 72h pi.: the nucleus contains a large replication compartment (RC) with numerous brilliant spots (red arrows) while fainter spots are visible in the nucleus outside the RC (white arrows) and in the cytoplasm (black arrows)(63X). All images acquired with a wide-field Zeiss Axiovert, Observer Z1, 1.4NA objective 63X.

## MATERIAL AND METHODS

### Viruses and Bacterial Artificial Chromosomes (BAC)

The TB40/E HCMV strain was obtained from a throat wash of a bone marrow transplant recipient patient (24) and its genome was cloned as a BAC in E.coli by replacing the non-essential US2 to US7 viral genes with the BAC vector pEB1097 (24, 25). This construct was later modified by inserting the GFP gene under the control of the murine CMV Immediate Early promoter (mCMV-MIEP) in the vector sequence, providing the TB40-GFP BAC (E. Borst, personal communication). This BAC has been maintained, amplified and mutated in DH10B bacteria grown in LB broth supplemented with the appropriate antibiotics. For the production of viruses from BACs, BAC DNA was first purified from bacteria using the PureLink HiPure Plasmid DNA Purification Kit (InVitroGen) or the NucleoBond XTRA BAC Kit (Macherey Nagel) according to manufacturer’s specific instructions. DNA was then transfected in MRC5 permissive human lung fibroblasts with X-tremeGENE^TM^ HP or X-tremeGENE^TM^ 9 transfection Reagents (Roche) following provided recommendations. When cytopathic effects reached nearly 100% of the cells, the content of the flask was harvested, centrifuged for 10 minutes at 2000RPM to remove cell debris and the supernatant was centrifuged at 25000 RPM (106000g) for 45 minutes at 16°C in a SW32Ti rotor (Beckman) on a 20% sucrose cushion. Alternatively, after the first centrifugation, supernatant could also be centrifuged at 20000 RPM (48500g) for 90 minutes at 16°C in a JA25.50 fixed angle rotor on a 20% sucrose cushion with similar virus yield. With both techniques, easily visible pellets were obtained under the cushion, resuspended in DMEM-20%FCS, aliquoted in vials and frozen at −80°C. The ANCHOR modified HCMV BACs were derived from the TB40-GFP BAC (a kind gift of Drs. E. Borst and M. Messerle). As a first proof of concept, the TB40-GFP BAC was initially modified by introducing an ANCH2 target sequence instead of the mCMV-MIEP-GFP gene. Briefly, the ANCH2 sequence and a kanamycine resistance gene were amplified respectively from the pUC18-ANCH2 (22) and the pORI6K-5FRT (a kind gift of Dr. M. Messerle) plasmids using PrimeStar Max 2X (TAKARA) according to manufacturer’s recommendations. The fragments were then purified, phosphorylated, ligated and the ligation product was used as template for a second amplification between new primers selecting the required product of ligation and introducing at both extremities 50bp homology sequences (H1 and H2) necessary for the final recombination of this product in the TB40-GFP BAC. This H1-Kana^R^-ANCH2-H2 cassette (Fig.2) has been cloned between the PvuII sites of the pGEM-7Zf(+) vector (PROMEGA) using NdeI or ApaI linkers (pGΔANCH2-kana). DH10B bacteria containing the TB40-GFP BAC were first transformed with the pKD46 vector encoding the arabinose inducible phage Red α, β and γ recombinases and then, with the purified H1-Kana^R^-ANCH2-H2 cassette. Recombinant clones were obtained, analyzed by BamHI digestion profile and the clone TB40-ANCH2-Kana was finally verified by sequencing (Fig.2). This BAC was amplified, purified and transfected into MRC5 human fibroblasts. Complete cytopathic effects were observed 4 weeks later and at this time, the content of the flask was harvested and viruses purified as described above. We next created the TB40-ANCHOR3 BAC by replacing the mCMV-MIEP-GFP sequence of the TB40-GFP BAC with a cassette containing the ANCH3 target sequence, a chimerical gene encoding the OR3 protein fused to GFP under the control of a SV40 promoter and a kanamycine resistance gene as a selection marker. This cassette was derived from the previously obtained pGΔANCH2-kana plasmid of which the ANCH2 sequence had been removed by PmlI digestion and replaced with an ANCH3 sequence, providing construct pGΔANCH3-kana. The OR3 gene had already be cloned in the peGFPc1 vector (Clontech) and a SV40 promoter was inserted directly upstream of the GFP-OR3 gene providing the pSVGO3 plasmid. The pSV40GFP-OR3 cassette was excised from the pSVGO3 plasmid and inserted in the MluI/PvuII sites of the pGΔANCH3-kana plasmid, creating the pG7ΔKanaSVOR3GFPANCH3 vector. Finally, the cassette of interest with the following structure H1-kana-OR3GFP-pSV40-ANCH3-H2 was excised by NdeI digestion, agarose gel purified and used for recombination in TB40-GFP containing DH10B bacteria. Obtained clones were screened by BamHI, EcoRI or HindIII restriction profiles and one of them, showing the expected modification, was named TB40-ANCHOR3 and further confirmed by PCR and DNA sequencing (Fig.2). This clone was then amplified, purified and transfected into MRC5 human fibroblasts. Complete cytopathic effects were observed 5 weeks later. At this time, viruses were purified and stored as described above. To create a pure ANCHOR-modified viral stock from this first production, 10^4^ MRC5 cells plated in 96 well plates (imaging grade, Corning Cell Bind) were infected with 0.5 virus per well. Albeit more than half of the wells did not display infection, one isolated green fluorescent plaque could be recovered and used to infect 10 ^4^ fresh MRC5 cells. After cell lysis, the culture medium was transferred to two T175 flasks of MRC5 cells which were incubated until cytopathic effects were estimated maximum (12 weeks from plaque picking). At this time, the content of the flasks was harvested and viruses purified as described, providing a stock of 8.10^8^ TB40-ANCHOR3 viruses.

**Figure 2.**
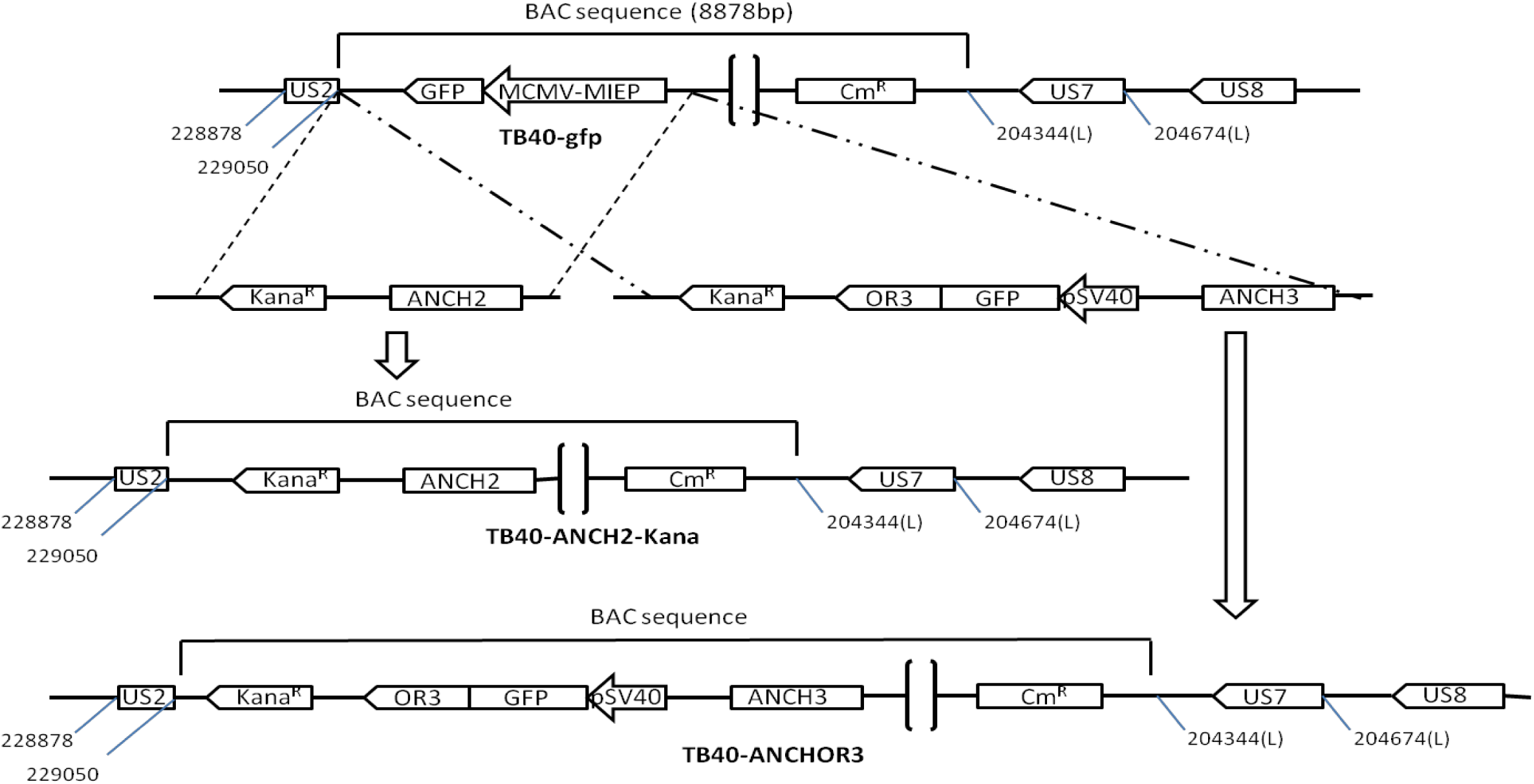
Construction of the ANCH2 or ANCHOR3 HCMV BAC used in this study and characterization of viruses derived from the TB40-ANCHOR3 HCMV BAC. Both ANCH2 and ANCHOR3 HCMV BAC were derived from the TB40-GFP BAC which contains the GFP gene under the control of a murine CMV immediate early promoter (pMCMV-MIEP-GFP) inserted in the vector backbone. This gene was replaced by the desired constructs without affecting any other viral gene or sequence. The TB40-ANCH2-Kana HCMV BAC displaying a single ANCH2 target sequence has been obtained by exchanging the MCMV-MIEP-GFP gene with an ANCH2-Kana^R^ cassette while the MCMV-MIEP-GFP gene has been exchanged by homologous recombination with a Kana^R^–OR3/GFPANCH3 cassette to create the TB40-ANCHOR3 HCMV BAC.

### Plaque forming assay

For quantification by plaque forming assay, 10^5^ MRC5 cells/well were plated in 24-well plates. The day after, culture medium is removed and cells are infected with various dilutions of the virus stock to be titrated. After two or sixteen hours contact, virus dilutions were removed and replaced with fresh supplemented culture medium containing 0.5% low-melting-point agarose and plaques were allowed to develop until they become visible. At this time, cells were fixed by addition of 1mL of 2% formaldehyde and plaques counted after staining with 0.02% methylene blue (26).

### Replication rate assay

To measure replication rates, 10^5^ MRC5 cells/well were seeded in two 24-well plates and infected with the TB40-GFP or the TB40-ANCHOR3 viruses at an MOI of 0.2. After 3 hours of contact, supernatants were removed, replaced with fresh medium and plates were incubated at 37°C under 5% CO2. One of the plates with a glass bottom was used to determine the number of cells and of fluorescent infected cells at each time point. Supernatants and cells were separately collected from duplicate wells of the second plate 1, 2, 4, 6, 8 or 10 days post infection. Supernatants were directly extracted with phenol and chloroform-isoamyl alcohol mixture and ethanol precipitated in the presence of HCMV-free carrier DNA. After centrifugation, DNA pellets were resuspended in 100µL sterile TE and conserved at 4°C. DNAs from the infected cells were purified using the NucleoSpin DNA RapidLyse kit (Macherey Nagel, ref. 740100.50) according to manufacturer’s instructions and eluted in a final volume of 200µL. Viral DNA quantification was performed in triplicate by qPCR using a LightCycler^R^480SYBR Green I Master kit (Roche, 04707516001). The cloned UL79 gene was chosen as the reference and the fragment flanked by primers UL79.1F (CAGATTAGCGAGAAGATGTCG) and UL79.1R (CAGGTTGTTCATGGTTTCGC) was amplified with a PCR efficiency ranging between 1.999 and 2.007.

### Cells, Culture, and Media

Human primary fibroblasts MRC5 (CCL-171) and human retinal pigmented epithelium cells ARPE-19 (CRL-2302), were obtained from ATCC and were grown in DMEM without phenol red (Gibco) supplemented with 10% FBS, 1x Penicilline-Streptomycine, 1 mM sodium pyruvate and 1× Glutamax (Gibco). Human umbilical vein endothelial cells (HUVEC, a kind gift of Dr. Melinda Benard) were grown in EGM-2 medium including supplement (PromoCell).

### Chromatin immunoprecipitation

ChIP assays were performed as described by Metivier *et al*. (27) with minor modifications. MRC5 cells infected at MOI 0.5 with TB40-GFP or TB40-ANCHOR3 viruses were treated 72h post-infection (pi.) with 1.5% formaldehyde for 10 min. Cross-link was stopped with 1M Glycin for 30 seconds and cells were washed with cold PBS. After nucleus preparation with Buffer NCPI (EDTA 10mM, EGTA 0.5mM, Hepes 10mM and TritonX-100 0.2%) and buffer NCPII (EDTA 1mM, EGTA 0.5mM, Hepes 10mM and NaCl 200mM) cell lysis was performed (10 mM EDTA, 50 mM Tris-HCl [pH 8.0], 1% SDS, 1× protease inhibitor cocktail [Roche Biochemicals, Mannheim, Germany]). Subsequently chromatin was either digested 3h at 37°C with DrdI and ScaI restriction enzymes and sonicated three times 10 seconds or not treated at all. Immunoprecipitations were performed overnight in the presence or not of 2 μg of selected antibody. Complexes were recovered by a 2 hr incubation with protein A Sepharose CL4B saturated with salmon sperm DNA. Beads were sequentially washed in buffer I (2 mM EDTA, 20 mM Tris-HCl, pH8.1, and 150 mM NaCl), buffer II (2 mM EDTA, 20 mM Tris-HCl, pH 8.1, 0.1% SDS, 1% Triton X-100, and 500 mM NaCl), buffer III (1 mM EDTA, 10 mM Tris-HCl, pH 8.1, 1% Nonidet P-40, 1% deoxycholate, and 250 mM LiCl), and three times with Tris-EDTA buffer. Washed resin was resuspended in elution buffer (1% SDS, 0.1 M NaHCO_3_) with 30-min incubation and the cross-link was reversed at 65°C overnight. DNA was purified with QIAquick columns (Qiagen, France). After immunoprecipitation with anti-GFP antibodies, PCR were performed with the following oligonucleotides: ANCH3-P7 (S) and (AS); CMV-P3 (S) CCGTACTTCGTCTGTCGTTT and (AS) TGTGTCTGTTTGATTCCCCG; CMV-P1 (S) ACGGCAAGTCCATAATCACC and (AS) GACCGATCCCACCAATTCTC; GFP-P1 (S) ACGTTGTGGCTGTTGTAGTT and (AS) GACTTCTTCAAGTCCGCCAT.

### High-Content Imaging

MRC5 cells are seeded at 10^4^ cells/well in Corning *Cellbind* black glass-bottom 96-well plates and infected twenty-four hours post-seeding with ANCHOR engineered HCMV at various MOI. For analysis, cells were directly stained with Hoechst 33342 (1μg/ml) and imaged using a Thermo Scientific Cellomics Arrayscan Vti microscope. Compartmental analysis was used to detect and quantify infection rate ie the number of GFP positive cells vs total number of cells. For spot counting and measure of viral DNA accumulation in the nuclei of infected cells, we used spot detector plugin for ImageJ with the following settings: spot radius 2, cutoff 0, percentile 7. Measurements were done for 1000 cells and average +/-SD were displayed from triplicates. Full analysis protocols for Arrayscan imaging are available upon request.

### IC50 calculation

In order to determine the IC50, a nonlinear regression was applied using the natural logarithm of the viral DNA content as dependent variable, the natural logarithm of the concentration as continuous predictor and the plate as covariate according to the following model:

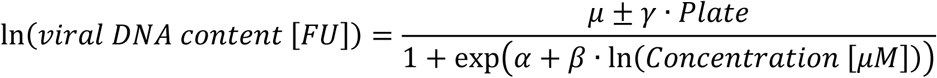

where α, β, γ and μ are the parameters of a non-linear regression.

The nonlinear regression was realized using R and the results are presented using the package ggplot2 (28).

### Fluorescence imaging

Live microscopy was performed using a Zeiss Axio Observer Z1, Apotome 2 wide-field fluorescence microscope. Conditions of acquisition are detailed in figure legends.

### Immunofluorescence

For immunofluorescence analysis, TB40-ANCHOR3 infected MRC5 cells were fixed in PBS containing 3.6% formaldehyde at different times pi., washed and then permeabilized in 10mM Hepes containing 0.5%Triton X-100 and 1% BSA. After washing, the cells were incubated for 1 hour at room temperature with the first antibodies diluted according to manufacturer’s recommendations, washed again and further incubated for 45 minutes in the appropriately diluted secondary antibodies. Cells were washed again and stained with Hoechst 33342 at the final concentration of 1µg/mL before fluorescence microscopic examination. For direct immunofluorescence of viral particles, diluted viruses were spotted on poly-L-Lysine pre-coated Ibidi glass bottom 35 mm dishes. Following a ten minute incubation, immunofluorescence was performed as described, with slight modifications. Due to the small size of viral particles, incubations were shortened to 30 min for primary and to 15 min for secondary antibodies.

### Correlative light and electron microscopy

MRC5 cells were grown on MatTek dishes with a finder grid (29) and infected with the TB40-ANCHOR3 virus at a MOI of 1. Four days post-infection, cells were fixed with 0.05% glutaraldehyde (GA) and 4% paraformaldehyde (PFA) for 30 min at room temperature. Image acquisition and analysis was performed on an Olympus IX81 epifluorescence microscope equipped with a x100 objective lens (UPlan SApo 1.4 oil), a SpectraX illumination system (Lumencore©) and a CMOS camera (Hamamatsu© ORCA-Flash 4.0). Stacks of 51 images each, with a step size of 0.1 µm, were taken. Cells were then fixed overnight with 2.5% GA in cacodylate buffer pH 7.2 and post fixed in 1% osmium in distilled water for 30 min. The samples were then rinsed in water and dehydrated in an ethanol series and flat embedded in Epon. Sections were cut on a Leica Ultracut microtome and ultra-thin sections were mounted on Formvar-coated slot copper grids. Finally, thin sections were stained with 1% uranyl acetate and lead citrate and examined with a transmission electron microscope (Jeol JEM-1400) at 80 kV. Images were acquired using a digital camera (Gatan Orius) at 200 and 2500X magnification. Alignments were performed as published (30).

## RESULTS

### Construction of the ANCHOR-HCMV BACs

Two ANCHOR-modified HCMV-BACs were derived from the TB40-GFP BAC (a kind gift of Drs. E. Borst and M. Messerle). The first one, TB40-ANCH2-Kana, was obtained by replacing the mCMV-MIEP-GFP gene of the TB40-GFP with a single ANCH2 target sequence (together with a selection kanamycine resistance gene). The TB40-ANCHOR3 BAC was constructed in a similar way but replacing the mCMV-MIEP-GFP sequence with a cassette containing the ANCH3 target sequence, a chimerical OR3-GFP gene driven by an SV40 promoter and a kanamycine resistance gene (Fig.2, see Material and Methods). Contrary to the TB40-ANCH2-Kana virus which requires separate transfection of an expression vector for its corresponding OR2 protein, the TB40-ANCHOR3 is autonomous in that it contains the ANCH3 target sequence and the gene encoding its cognate OR3 protein (fused to the GFP gene).

### Viruses derived from the TB40-ANCHOR-HCMV BACs are infectious

Transfections of the purified TB40-ANCHOR BACs in MRC5 fibroblasts were poorly efficient, likely due to their very large size and to low transfection efficacy of the cells. Only sparse foci of modified cells were observed 10 days post transfection. However, nearly 100% cytopathic effects were reached 4 to 5 weeks after transfection, indicating that the rare cells which were transfected with the TB40-ANCHOR BACs produced viruses that were fully infectious. In order to confirm infectivity and to assess whether TB40-ANCHOR3 viruses had conserved the epithelial and endothelial tropism of the original TB40 HCMV strain, TB40-ANCHOR3 viruses were used to infect new MRC5, HUVEC or ARPE-19 cells, respectively fibroblasts, endothelial and retinal epithelial cells. The three cell types became readily infected with appearance of fluorescence and fluorescent spots several hours pi, confirming infectivity of the ANCHOR engineered viruses and the preservation of the original cellular tropism (Fig. 1b, Supplementary Fig.S1). However, infection efficacy and kinetics were clearly cell type dependent as the number of infected cells in the presence of the same amount of virus, varied largely from one cell type to the other. As a first estimate of its replication capacity, the TB40-ANCHOR3 viral stock was titrated using two different techniques: a fluorescence assay and a classical plaque forming assay. MRC5 cells were infected with different dilutions of the viral stock for 2 or 18 hours, washed and incubated as described. The plate of the fluorescent assay was analyzed with an automated ArrayScan microscope 60h pi., while the plate with the plaque forming assay was maintained at 37°C for 12 days and then fixed, stained and analyzed. As shown in Table 1, very similar results were obtained with both techniques indicating that the viruses infecting the cells render them fluorescent and are also able to induce a complete lytic cycle. Interestingly, the contact time between cells and viruses matters as the longer infection times systematically result in higher titers than the shorter ones, whichever the technique is used. However, even if both techniques are reliable, the fluorescent one is clearly less labor-intensive, more rapid, robust and reproducible and was hence adopted for all further titrations

**Table 1.** Measure of the TB40-ANCHOR3 HCMV stock virus titer. Titer was measured using a classical plaque forming assay and a fluorescence assay. See Material and Methods for details. Whatever the technique, measured titers are significantly higher when infection time is increased from 2h to 18 h. However, very similar results are obtained with both techniques.

| Measured virus titer |  |  |  |  |
| --- | --- | --- | --- | --- |
| *μL stock virus* | Plaque-forming assay |  | Fluorescence |  |
|  | *2h infection* | *18h infection* | *2h infection* | *18h infection* |
| $10^{-4}$ | $6,0 \cdot 10^7$ | $2,0 \cdot 10^8$ | | |
| $10^{-3}$ | $5,2 \cdot 10^7$ | $1,6 \cdot 10^8$ | $2,1 \cdot 10^7$ | $1,2 \cdot 10^8$ |
| $10^{-2}$ | $2,8 \cdot 10^7$ | | $3,9 \cdot 10^7$ | $4,2 \cdot 10^8$ |
| $10^{-1}$ | | | $1,2 \cdot 10^8$ | $1,9 \cdot 10^8$ |

### ANCHOR modification does not interfere with virus replication rate

To quantify more precisely data about their replication kinetics, we infected MRC5 cells with TB40-GFP and TB40-ANCHOR3 viruses at an MOI of 0.2, and measured the number of viral genomes present in cells and in their supernatants until 10 days post infection (Table 2). Values measured in supernatants were compared with those published for the TB40-BAC4 (31) and are presented in Fig.3. Both in supernatants and in cells, the number of TB40-ANCHOR3 genomes is larger than the one determined for TB40-GFP, suggesting that TB40-ANCHOR3 replicates more efficiently. TB40-GFP and the TB40-BAC4 strains produce similar number of genomes in supernatants. These results show that the presence of the ANCHOR sequences in the viral genome does not impair its replication nor induces important functional deletion.

**Table 2.** Replication kinetics of TB40-GFP and TB40-ANCHOR3 viruses. MRC5 cells were infected at an MOI of 0.2. Cells and supernatants were harvested on days 2, 4, 6, 8 and 10 post-infection and DNA purified from each sample. Total numbers of viral genomes were determined for each sample using qPCR. Cells and infected cells were counted in a parallel plate. Each measure was made in triplicate and mean values of the two wells per time point are given for each day (except for day 10 pi.).

| Table 2 |  |  |  |  |  |  |
| --- | --- | --- | --- | --- | --- | --- |
| Day pi. | TB40-GFP |  |  | TB40-ANCHOR3 |  |  |
| | Cells/well($\times 10^{-3}$) | Infected cells/well($\times 10^{-3}$) | Viral genomes /infected cell($\times 10^{-3}$) | Cells/well($\times 10^{-3}$) | Infected cells/well($\times 10^{-3}$) | Viral genomes /infected cell($\times 10^{-3}$) |
| 2 | 129/132 | 9.1/9 | 0.17/0.13 | 111/114 | 9.6/13.9 | 9.4/8.3 |
| 4 | 316/295 | 26.7/40.6 | 0.21/0.44 | 254/288 | 27.3/33.2 | 10.7/16.5 |
| 6 | 282/289 | 76.4/78.8 | 0.71/0.83 | 226/256 | 50.5/36.8 | 14.3/19.2 |
| 8 | 232/240 | 79.4/79.3 | 1.3/1.4 | 135/210 | 66.1/44.5 | 9.4/12.7 |
| 10 | 234 | 89.6 | 0.54 | 167 | 89.1 | 2.3 |

**Figure 3.**
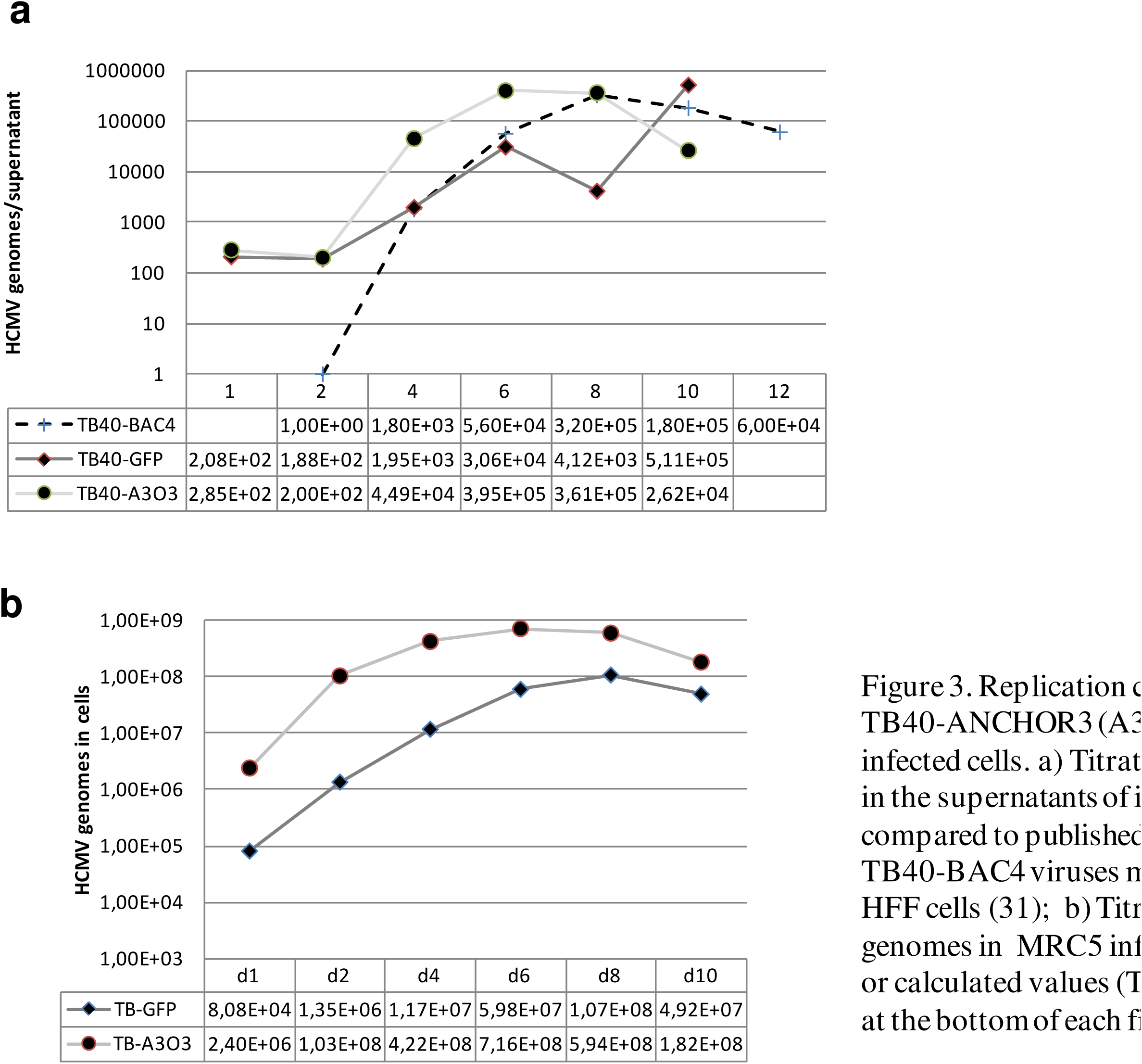
Replication curves of TB40-GFP or TB40-ANCHOR3 viruses in MRC5 infected cells. a) Titration of HCMV genomes in the supernatants of infected cells, compared to published growth kinetics of TB40-BAC4 viruses measured by TCID50 in HFF cells (31);b)Titration of HCMV genomes in MRC5 infected cells. Measured or calculated values (TB40-BAC4) are given at the bottom of each figure.

### Viruses derived from the TB40-ANCHOR-HCMV BAC are fluorescent and mature

TB40-ANCHOR3 viruses from the purified viral stock were immobilized on poly-lysine treated glass slides and stained with Hoechst and anti-pp28 or anti-gB antibodies. As shown in Fig.4, Hoechst, GFP, anti-pp28 and anti-gB signals perfectly superimpose suggesting that these viral particles are mature, tegumented, enveloped and contain DNA and OR3-GFP proteins.

**Figure 4.**
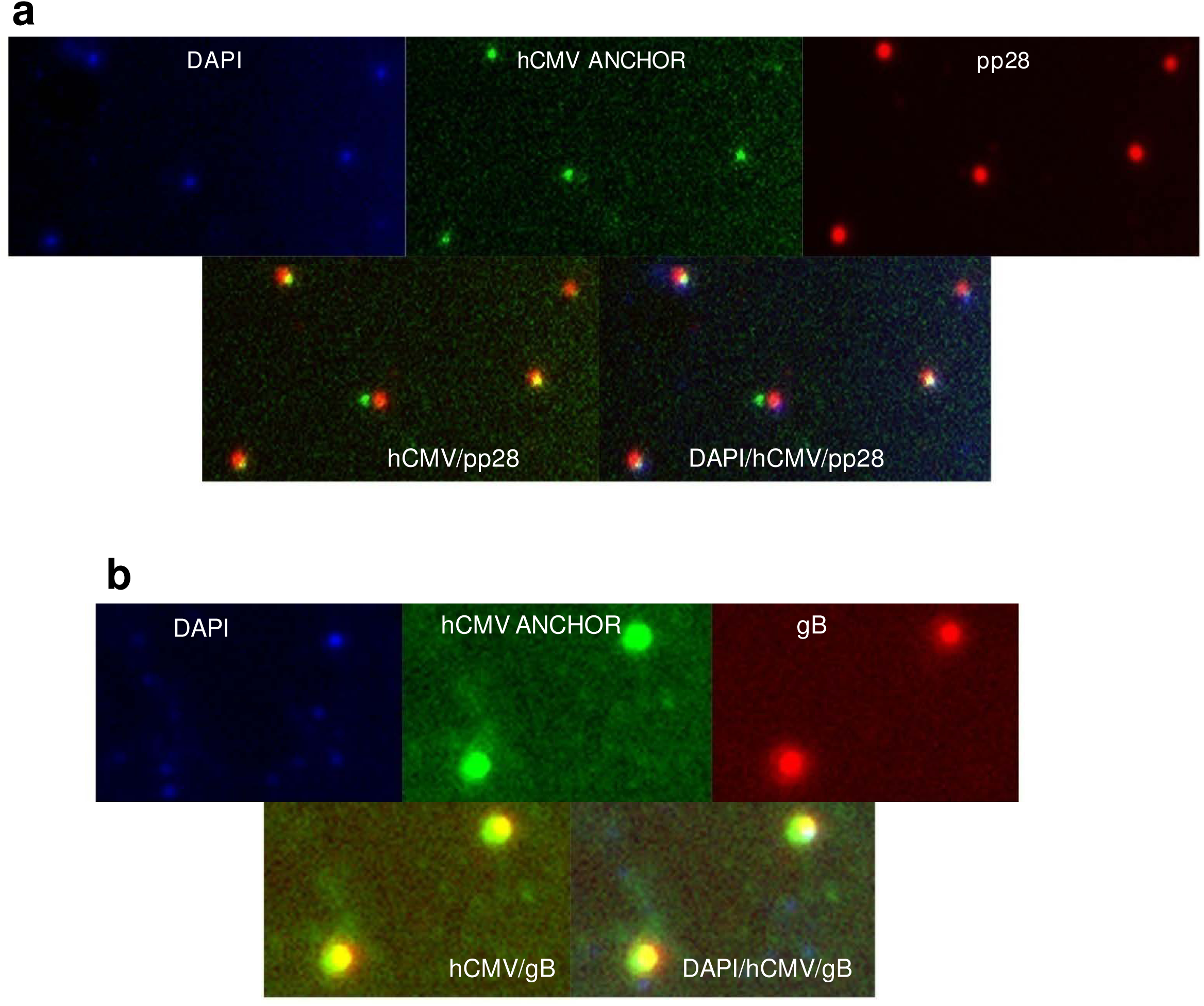
Characterization of TB40-ANCHOR3 viruses. Viral particles derived from the TB40-ANCHOR3 HCMV BAC contain DNA, OR3-GFP proteins and stain for pp28 tegument proteins (a) and for gB envelope proteins (b) and therefore likely correspond to mature viruses, possibly with OR proteins bound to the encapsidated genomes. Images acquired with a wide-field Zeiss Axiovert, Observer Z1, 1.4NA objective 63X

### OR-GFP proteins effectively bind to ANCH sequences in ANCHOR-HCMV infected cells

Binding of OR proteins to cognate engineered ANCH target sequences was previously demonstrated in pro-and eukaryotic systems (22). We confirmed that the same held true in cells infected with our HCMV-ANCHOR. For this purpose, chromatin immunoprecipitation (ChIP) experiments were performed on cells infected with TB40-GFP or TB40-ANCHOR3 viruses using antibodies against GFP. As shown in Fig. 5a, DNA immunoprecipitated from TB40-ANCHOR3 infected cells is strongly enriched in ANCH3 sequences, confirming OR3-GFP proteins bind to the ANCH3 target sequence. Enrichment in the adjacent GFP sequence suggests spreading of OR3-GFP onto neighboring DNA. No significant enrichment of more distant sequences was observed. Similarly, no significant enrichment was observed in DNA from cells infected with the TB40-GFP virus immunoprecipitated with anti-GFP antibodies.

**Figure 5.**
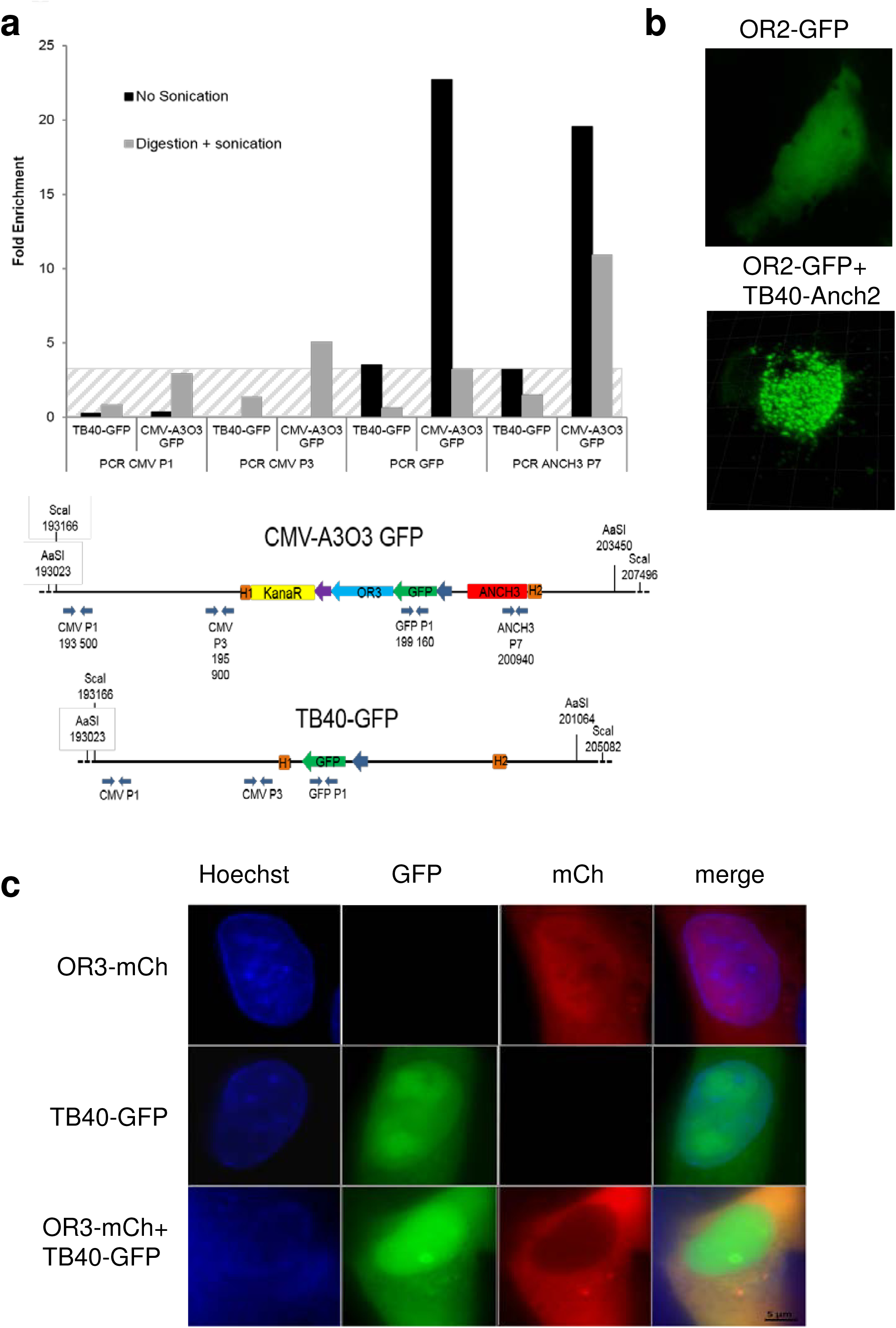
OR-FP specifically bind to their target ANCH sequence generating the spots observed in fluorescence microcospy. a)ChIP experiment showing that Chromatin extracted from TB40-
ANCHOR3 HCMV infected MRC5 cells (but not from TB40-GFP Infected cells) and immunoprecipited with anti-GFP antibody is only enriched in ANCH3 and GFP sequences; hatched section corresponds to background noise, since no ANCH3 sequence is present in the TB40-GFP virus; b) MRC5 cells were transfected with a single vector expressing OR2-GFP and then infected or not with TB40-ANCH2-Kana viruses; spots appear only when OR-FP and the corresponding ANCH target sequences on the viral genomes are present simultaneously in the same cell. Images aquired 24 h post transfection and 72h post infection with a wide-field Zeiss Axiovert, Observer Z1, 1.4NA objective 63X or a Zeiss LSM 510 NLO; c) MRC5 cells were transfected with an expression vector for OR3-mCherry or infected with TB40-GFP or transfected and infected simultaneously; no spot are observed in any situation indicating that nor OR3 proteins neither TB40-GFP genomes form unspecific spots, even when present simultaneously in the same cell.

### In infected cells, fluorescent spots result from OR-FP binding to ANCH-HCMV genomes

To determine the nature of the observed spots (Fig.1b, Supplementary Fig.1), we took advantage of our TB40-ANCH2-Kana virus which contains a single ANCH2 sequence but no OR-FP gene. When MRC5 cells were solely infected with the TB40-ANCH2-Kana viruses, no fluorescence was observed (results not shown). Transfection of an expression vector for OR2-GFP proteins in uninfected MRC5 cells resulted in uniform fluorescence in all cell compartments while infection of OR2-GFP transfected cells with the TB40-ANCH2-Kana virus resulted in the appearance of numerous bright spots 72 hours post-infection (Fig.5b). As a control, MRC5 cells which had been transfected with an expression vector for OR3-mcherry were infected with the TB40-GFP virus: as shown in Fig5c., 72hours pi., doubly fluorescent (red and green) cells did not display any spot similar to those observed in Fig. 5b or S1 despite the fact that a nuclear structure resembling a replication compartment is clearly visible (Fig.5c). These two experiments together demonstrate, on one hand, that OR-FP proteins or viral genomes alone do not form any spot and, on the other hand that OR-FP proteins do not form non-specific spots on HCMV genomes. Therefore, taken together with the ChIP experiments, these results demonstrate that ANCHOR-HCMV fluorescent spots result from the specific accumulation of OR-FP proteins on the corresponding ANCH sequences inserted in viral genomes.

### The ANCHOR cassette is stable in the recombinant virus

We have tested the stability of the ANCHOR phenotype after massive amplification from a single TB40-ANCHOR3 virus up to a stock of 8.10^8^ infectious particles. Viruses of this stock were used to infect MRC5 cells which were fixed and stained with various anti-HCMV antibodies at different times post-infection. We found that >90% of pp28 (tegument) positive cells were also positive for OR-GFP revealing that despite an amplification factor of nearly 10^9^, less than 10% of the final viruses had lost the ANCHOR phenotype (Results not shown). It is noteworthy that a similar situation was also observed for the TB40-GFP stock in which only 95% of the viruses were positive for both UL44 and GFP (Results not shown).

### Real-time visualization of ANCHOR -HCMV infection in living human cells

TB40-ANCHOR3 viruses were used to infect MRC5 fibroblast for time-lapse imaging of infection progression in live cells. Diffuse GFP fluorescence attributable to the OR3-GFP proteins was first detected in the cytoplasm as well as in the nucleus of the infected cells between 4 and 5 hours pi. This duration likely corresponds to the time required for the virus to attach and enter a cell, travel to the nucleus and to express its first genes. Interestingly, during the same lapse of time, infected cells transiently round out before recovering their usual spindle shape (Supplementary fig. S2). About 16 hours after infection, faint spots can be detected in the cells’ nuclei (Fig.6a). The number of spots increases during the two or three following days (Fig.6b), but these remain confined in small peculiar areas (Fig.6b) which fuse at the end (Fig.6c, Supp.Video1). About 72h to 95h pi., depending on cells, a single, large and well demarcated area containing up to several hundreds of intense spots occupies most of the nuclear space. This nuclear area is highly reminiscent of the replication compartment (RC) which has previously been associated with CMV intranuclear inclusion bodies (32) and later defined as the site of viral DNA replication and replication specific protein accumulation (33, 34). To better characterize this specialized area, we performed immunofluorescence staining of ANCHOR3-HCMV infected cells with anti-UL44 antibodies. UL44 encodes the polymerase associated processivity factor which is described to be specifically recruited to RC (34). Results presented in Fig. 7 clearly show that this well demarcated nuclear area containing most of the spots (and the most intense ones) is also precisely co-stained with the anti-UL44 antibody, indicating it is indeed the RC. Interestingly, UL44 distribution clearly evolves during the course of infection but always superimposes to the areas where HCMV genomes are also observed. This is true at 24 hpi., when viral genomes are still moderately amplified and present in limited, rather small areas (Fig.7a) but also at 72hpi., when the different replication zones have fused in a large replication compartment occupying most of the nucleus (Fig.7b). In addition to the most intense spots observed in the RC, numerous fainter spots are clearly visible in the rest of the nucleus and in the cytoplasm (Fig.6d) where they are especially abundant in a large demarcated, rounded region adjacent to the nucleus 72h pi. (Fig.7b). As it is now largely admitted that viral tegument and envelope are acquired in a specialized cytoplasmic compartment, we tried to better define the zone containing these cytoplasmic spots by IF staining for tegument (pp28) and envelope (gB) viral proteins. In our context of viral infection, pp28 presents a punctuate distribution in the whole cell early after infection (Fig.7c) but after 72h., few pp28 remains in the nucleus while most of it accumulates in a juxtanuclear region as described by others (Fig.7d) (35). Interestingly, at this time, numerous faint spots are also present in the very same zone. The same holds true for gB staining 72hpi. which accumulates in a similar domain where numerous HCMV spots are also clearly visible (Fig.7e). It is therefore very likely that this structure is the Assembly Compartment which overlaps the Endoplasmic Reticulum-Golgi-Intermediate Compartment (ERGIC) where naked capsids acquire their tegument and envelope (36, 37). Interestingly, the HCMV genomes remain well visible in addition to the IF targeted proteins, indicating that enough GFP protein from the ANCHOR system survive the immune-fluorescence procedure, in particular fixation, and that the two approaches are therefore compatible allowing analysis and co-localization of viruses or viral genomes with any cellular or viral protein of interest. Following the formation of the large unique RC, no appreciable change in viral accumulation or cellular morphology seems to occur for several hours. However, after this apparent quiescence or lag period, membrane rearrangements and cytoplasmic “bubbling” appear at one or both poles of the cell. These events rapidly amplify and suddenly result in cell fragmentation and death, similar to “blebbing” (38), only leaving fluorescent scraps (Supplemental video 2 & 3). It is noteworthy that each cell presents its own infection time course and some cells undergo a complete cycle from fluorescence appearance to “blebbing” and fragmentation in less time than the lag period between the mature RC and the cytoplasmic “bubbling” of some others (Supplemental video 4).

**Figure 6.**
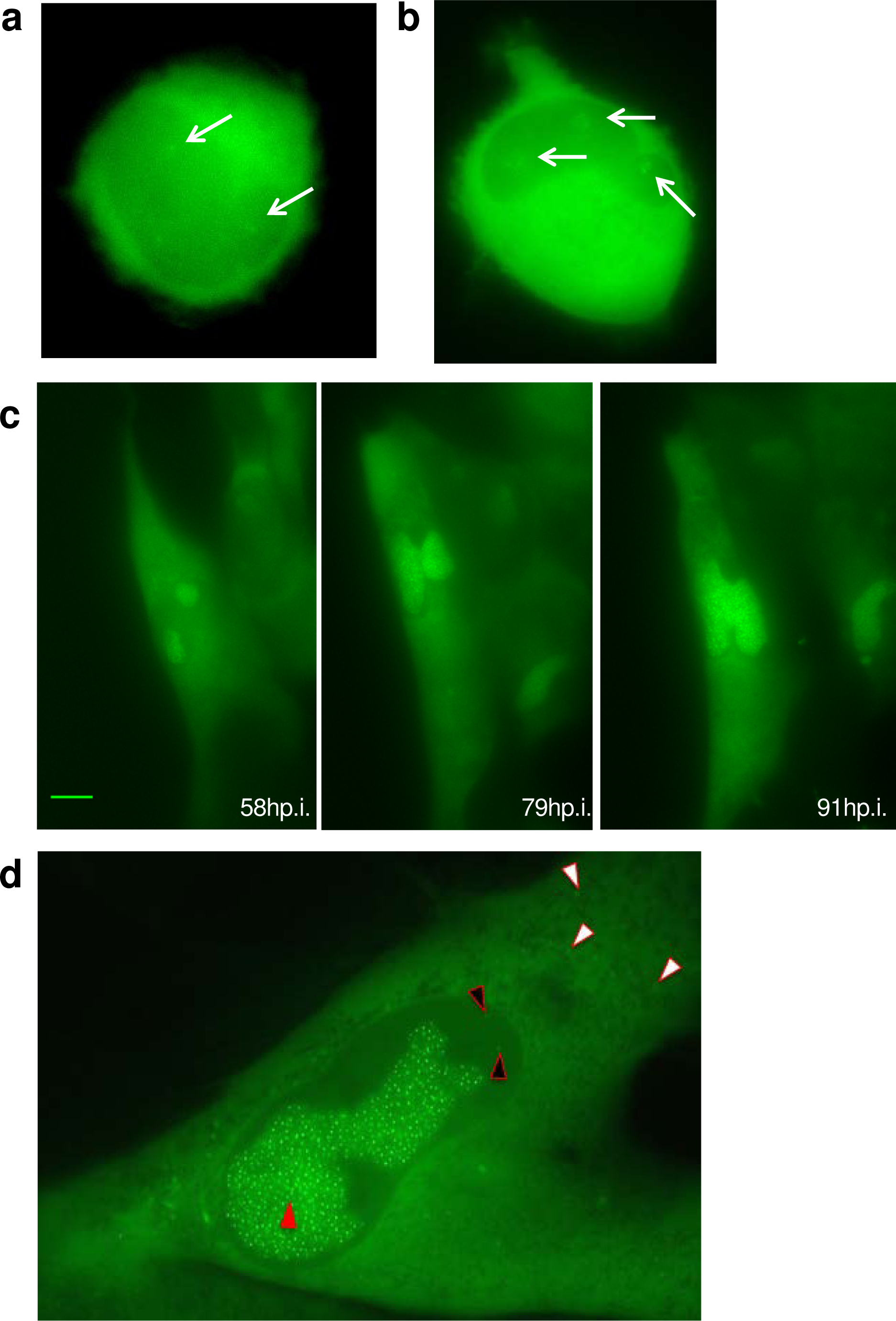
Visualization of ANCHOR-HCMV infection steps in living cells. MRC5 cells were infected with TB40-ANCHOR3 HCMV viruses at an MOI of 0.5; a) about 16-17 hours pi, some very faint spots appear in infected cells which possibly correspond to incoming viral genomes (white arrows, 63X); b) distinct areas suggestive of pre-replicative sites develop around the initial spots which multiply while these areas increase in size (white arrows, 63X); c) later in infection (around 70-80h pi.), these areas fuse in a unique putative replication compartment (RC) which continues to grow (40X, scale bar 10μm); d) an infected MRC5 cell imaged 72h pi.: the nucleus contains a large replication compartment (RC) with numerous brilliant spots (red triangles) while fainter spots are visible in the nucleus outside the RC (black triangles) and in the cytoplasm (white triangles)(63X). All images acquired with a wide-field Zeiss Axiovert, Observer Z1, 1.4NA objective 40X or 63X.

**Figure 7.**
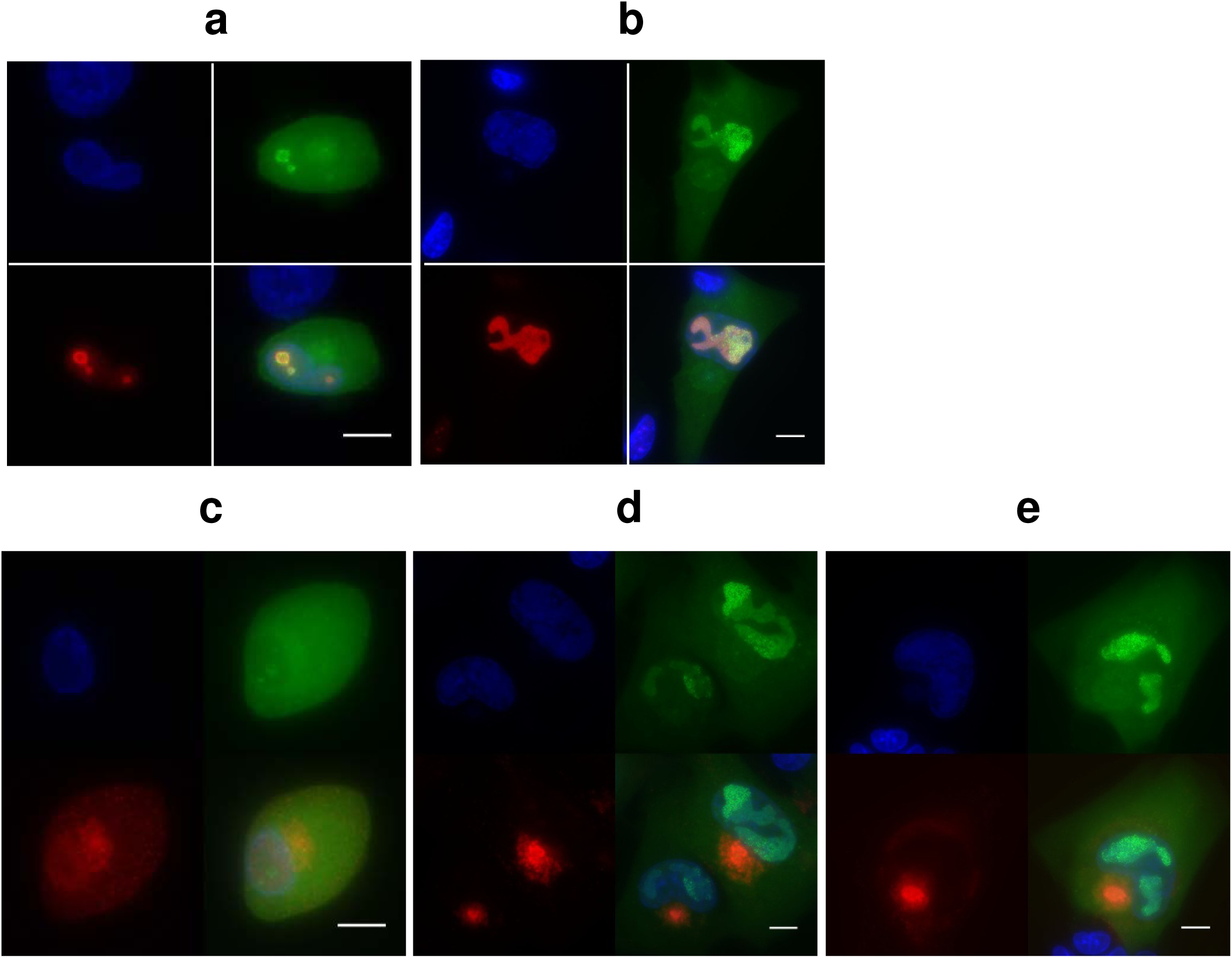
The putative replication compartment stains for the polymerase associated processivity factor pUL44 while pp28 and gB proteins accumulate in a region close to the nucleus suggestive of the Assembly Compartment. TB40-ANCHOR3 HCMV infected MRC5 cells were stained for pUL44 at different times pi. a) 24h pi. b) At 72h pi., only the putative RC is positive for pUL44 confirming its status. For each time point, upper left: Hoechst 33342, upper right: OR-GFP fluorescence, lower left: anti-pUL44, lower right: merge. TB40-ANCHOR3 HCMV infected MRC5 cells were stained for the pp28 tegument protein at different times pi. (c and d) or for the envelope gB protein (e). c) 24h pi., pp28 is already expressed but appears diffuse in the whole cell. d) On the contrary, 72h pi., pp28 is concentrated in a large region close to the nucleus where OR-GFP spots are also visible suggesting this region is the Assembly Compartment. For each time point, upper left: Hoechst 33342, upper right: ORGFP fluorescence, lower left: anti-pp28, lower right: merge. e) TB40-ANCHOR3 HCMV infected MRC5 cells were stained for the gB envelope protein 72h pi. showing accumulation in a region close to the nucleus, especially in the central area of this region; upper left: Hoechst 33342, upper right: OR-GFP fluorescence, lower left: anti-gB, lower right: merge

### Replicating viral genomes associate with preformed capsids

It is generally accepted that HV replicating genomes associate in the nucleus with preformed capsids of which the TER complex encoded by UL89, UL56 and UL51 captures and internalizes viral DNA through the portal complex (UL104) (39, 40). We analyzed cells at this stage of infection by correlative fluorescence/electron microscopy (Fig.8a). Electron microscopy revealed different types of capsids (Fig.8b), reminiscent of previously described A, B and C forms (41) but also possibly other forms (Fig.8c). When merging fluorescence and electron microscopy images, fluorescent spots and viral capsids nicely superimposed at the periphery of the RC (Fig.8d). The chosen area (yellow square in Fig.8a) shows four capsids on the electron-micrograph (Fig.8d), three containing material (type B?) and an empty looking one. Interestingly, the fluorescence staining type B capsids (which are at the edge of the RC) is weaker than the one associated with the other capsid, but equivalent between type B capsids, suggesting that these capsids already contain a single viral genome. On the other hand, the empty capsid could be linked to a replicative structure containing more than one viral genome.

**Figure 8.**
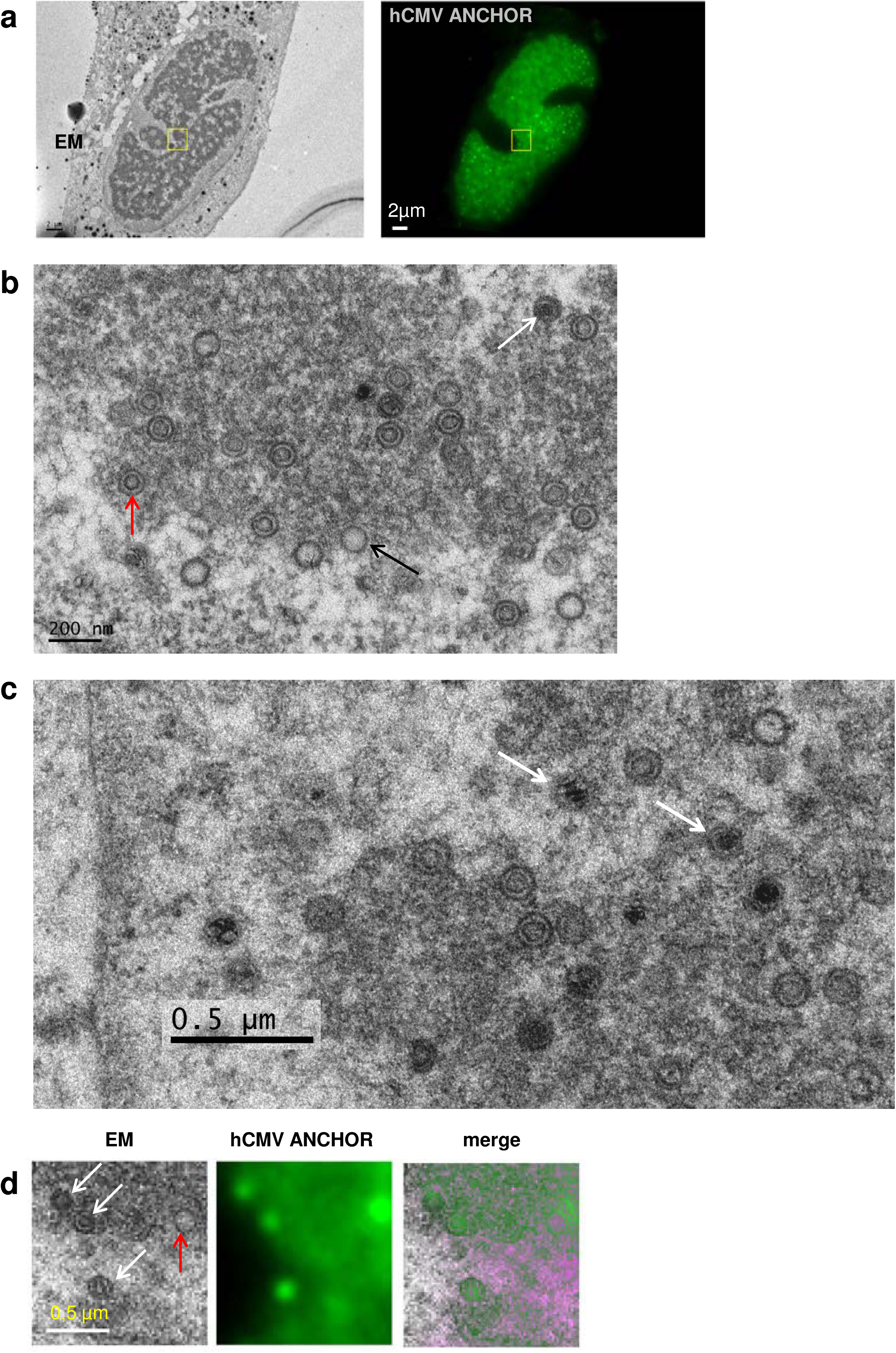
Visualization of infected cell by correlative fluorescence-electron microscopy. a)Cells infected with TB40-ANCHOR3 HCMV were first analyzed by fluorescence microscopy 96h pi. and then fixed and processed for electron microscopy examination (scale bar 2μm). The zone delimited by a yellow square has been further analyzed with higher magnification in d); b) in the RC, electron microscopy reveals different forms of capsids, possibly the type A, B and C capsids shown by black, red and white arrows respectively (scale bar 200nm); c) at higher magnification, other, possibly more diverse, forms of capsids seem to appear, with some containing dense fragmented material (white arrows) d) in the chosen area (corresponding to yellow box in a), 4 capsids are observed by EM, which all correspond to fluorescent spots. Three appear as B forms (white arrows) and one as a A form (red arrow). Spots associated with the three type B capsids present similarly weak intensities suggesting they contain a single viral genome. On the contrary, the type A capsid coincides with a much brighter spot and could be associated to a replicating structure with several genomes (scale bar 500nm). Pink color is an artefact resulting from the merging procedure.

### The ANCHOR technology enables quantitative tracking of HCMV infection

The number of ANCHOR spots present in a peculiar cell can be determined using the Image J particle detector software (Particle detector & tracker). In the cell illustrated in Fig.9, we detected n=1155 ANCHOR foci, distributed between the RC (n=1005), the remaining non-RC nucleoplasm (n=16) and the cytoplasm (n=134). Fluorescence intensity of the observed spots was highly variable (Fig.9a and b) and could be quantified (Fig. 9c) using an approach similar to the one that enabled precise quantification of E.coli replisomes and yeast telomerase (42). Briefly, we used the 3D interactive surface plot plugin for ImageJ that converts fluorescence intensity into arbitrary fluorescence units in the Z axis of the 3D reconstruction. Therefore, foci are not represented as 2D dots but as 3D peaks where the Z values correspond to fluorescence intensities. The diffuse background GFP fluorescence in the cytoplasm and the nucleus (outside the RC) appears in dark blue in Fig.9c and may be assigned a value of 90 arbitrary units (AU) on the color scale of Fig.9. In the same areas, pale blue regions and spots corresponding to +/-120AU, are also visible with some of the peaks superimposing with the spots present on Fig 9a and b. Interestingly, the RC itself is delimited by a line of spots of the same color. Inside the RC, colors are not distributed along a uniform gradient but rather in well defined successive, concentric zones of which the mean intensity radially decreases from the center (containing 240 and 210AU spots) to the periphery and which are separated by +/-30AU (Fig.9c and d). A similar distribution is also observed in ARPE-19 cells (Fig.S3) suggesting it results from a phenomenon common to all infected cells.

**Figure. 9.**
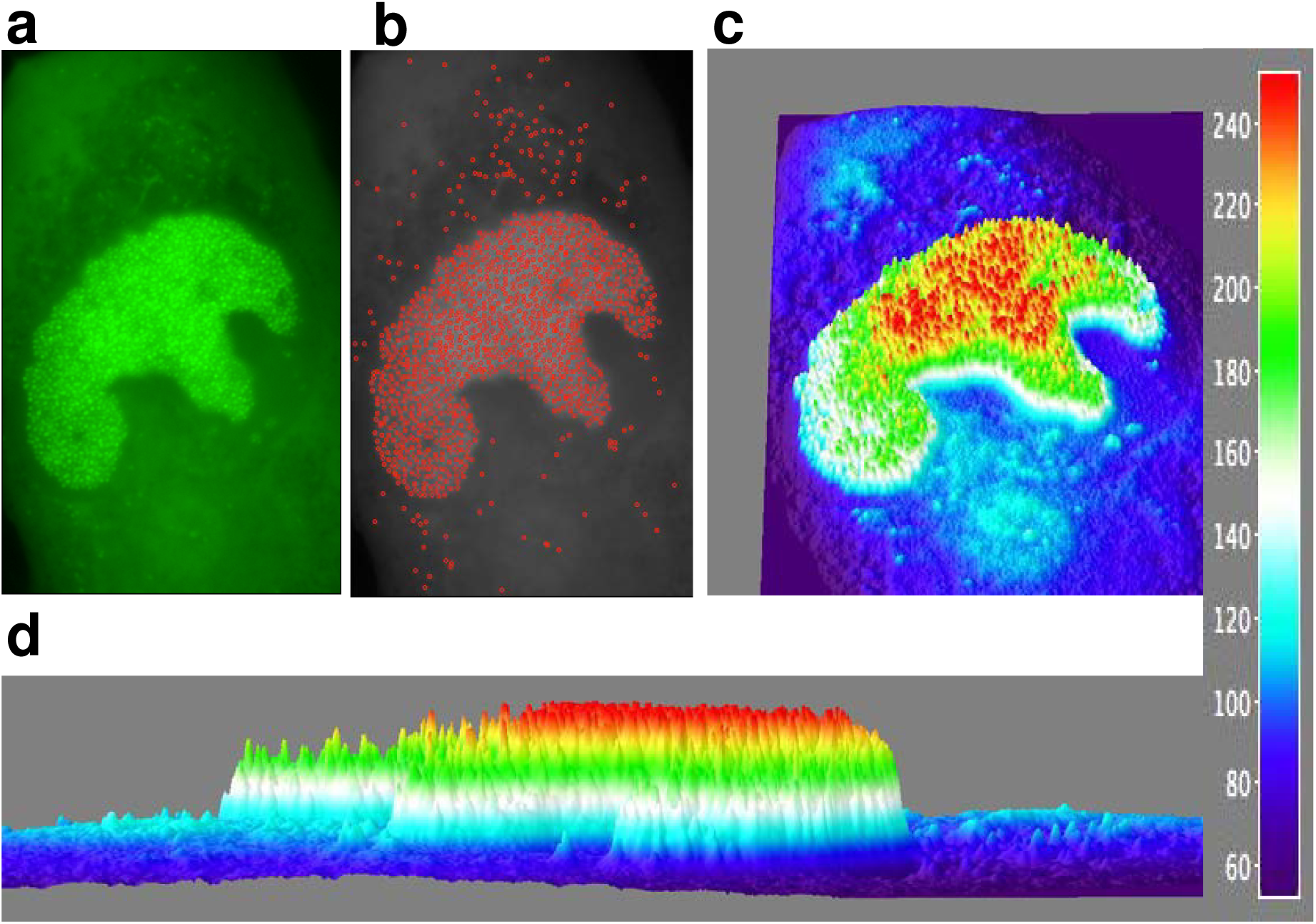
ANCHOR technology allows quantification of HCMV infection in live cells. a)Fluorescent spots in an MRC5 cell infected with TB40-ANCHOR3 HCMV 72h pi.; as demonstrated above, each spot corresponds to one or several fused genomes; b) same image as a) but treated to filter out the fluorescence background and then processed using spot detector plugin for ImageJ (spot radius 2, cutoff 0, percentile 7) allowing detection of 1155 ANCHOR foci: 1005 particles in the RC, 16 nuclear particles outside of RC and 134 cytoplasmic particles; c) images were then converted into 3D intensity surface plot (perspective) or, d) into a picture in X,Z to assess particles intensity. A single viral genome corresponds to 30 arbitrary fluorescence units (FU). In the RC, all spots harbor between 2 and 5 viral genomes and this number decreases from the center to the periphery, suggesting a highly organized territory. Outside the RC, only unique genomes are observed (see text for explanations).

### ANCHOR-HCMV is a new tool for rapid and cost-effective assessment of anti-viral compounds

To date, quantitative information about the presence of HCMV or its replication in a sample mainly relies on qPCR based technologies which, despite being very tedious, remain indispensable from a clinical point of view (43). For this kind of investigation and biotechnological applications, the ANCHOR technology also appears as a very promising alternative. We have analyzed infection kinetics of MRC5 cells in the presence or not of Ganciclovir, a compound widely used to treat HCMV infection (44, 45, 46). We have developed an Array-scan based custom algorithm for automated image analysis. To quantify the viral DNA content of cells infected with our TB40-ANCHOR3 HCMV viruses, this algorithm is remarkably efficient and enables direct determination of infection and replication rates per cell and/or per population (Fig.10a). Imaging of Ganciclovir treated cells revealed drastically reduced viral DNA content. Despite initial appearance of fluorescent particles in discrete nuclear domains, subsequent massive amplification and RC formation were inhibited (Fig.S4). Using various concentrations of Ganciclovir, IC50 was determined to be 2.26 µM, a value within the range measured by other techniques while IC90 was measured to be 8.435µM (47). When two independent plates were analyzed, intra-and inter-plate variability was remarkably low with a correlation coefficient of 0,97 (Fig.10b) demonstrating that our experimental approach is highly reproducible and robust. We next tested infectivity and response to drug treatment in parallel on two different cell lines: in a pilot experiment, we found that at an MOI of 0.5, the infection rate of MRC5 cells increased from 15 to 75% between 24 hours and 10 days p.i. In the presence of 2.5µM Ganciclovir, this increase was limited to 40% between days 7 and 10, with no effect at 24h p.i., as expected for a drug blocking the viral polymerase and not the virus entry. In contrast, the infection rate of ARPE-19 cells infected at an MOI of 0.5 remained constant between 1 and 3% during the entire experiment, with no evident effect of Ganciclovir (Fig.10c). Viral DNA content was reduced by 95-100% in Ganciclovir treated MRC5 cells 7 or 10 days pi. while surprisingly, in ARPE-19 cells, the viral DNA content increased until day 7 pi. Despite a slight decrease at day 10 p.i., no effect of Ganciclovir was observed at 2.5µM (Fig.10d) in these cells which required 12.5µM to completely abolish HCMV replication, indicating that sensitivity to Ganciclovir is cell type dependent (data not shown). The ease of this quantification paves the way for innovative screening strategies in the search of new anti-viral drugs.

**Figure 10.**
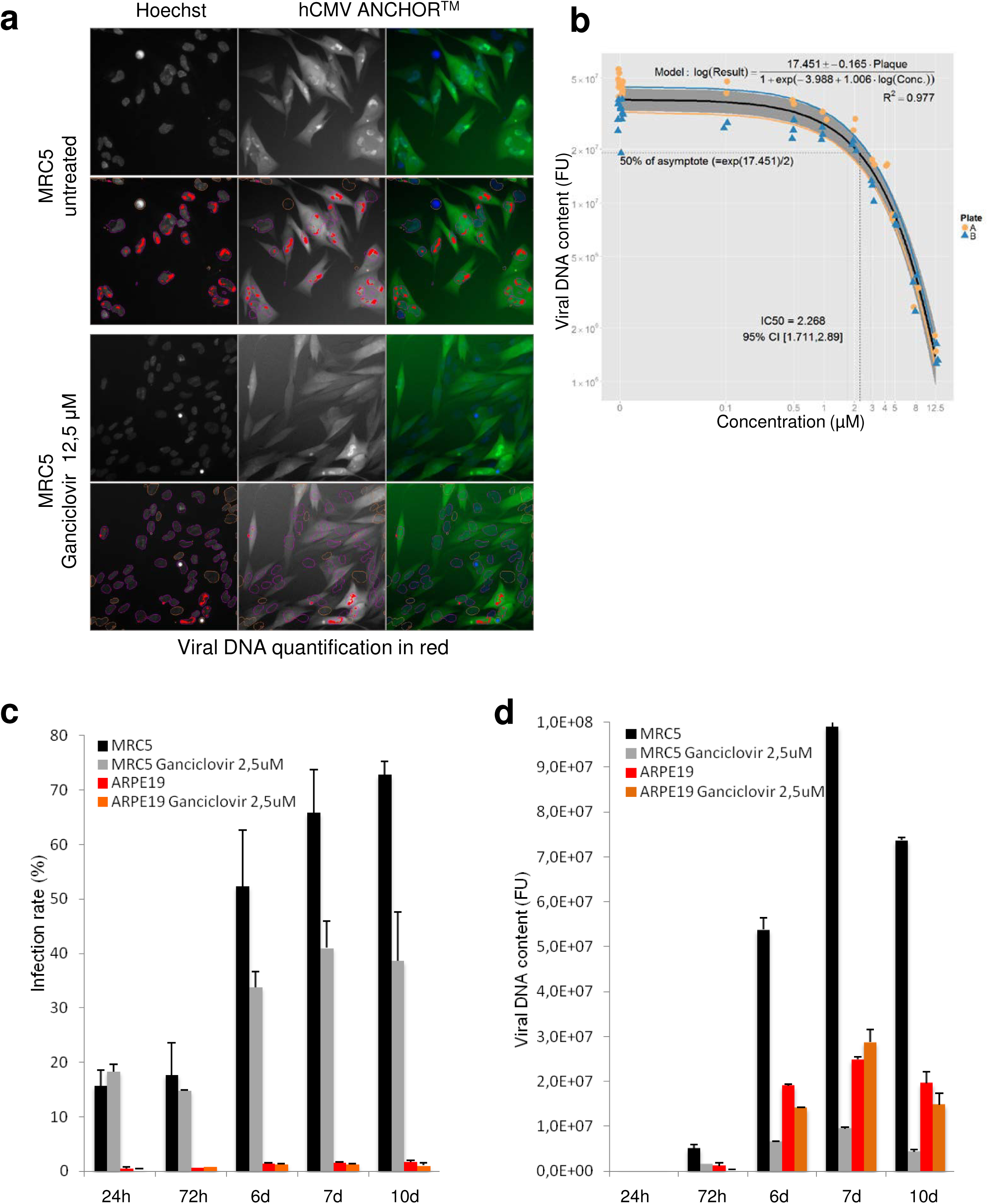
Cell type specific effect of Ganciclovir on TB40-ANCHOR3 HCMV infection and IC50 determination by automated high content imaging. a) TB40-ANCHOR3 HCMV infected cells were treated or not with 12.5μM Ganciclovir. 72 hours pi, cells were observed using a Thermo Scientific Cellomics Arrayscan Vti microscope and images analyzed with the compartmental analysis algorithm allowing detection of viral DNA (in red, see Material and Methods for explanation); b) similar experiment as a) but TB40-ANCHOR3 HCMV infected cells were treated with increasing doses of Ganciclovir. Results of the quantification were plotted against Ganciclovir concentrations (see text) allowing precise determination of IC50 and IC90. Two identical experiments were performed on separate plates (A and B); c) time course of TB40-ANCHOR3 HCMV infections in MRC5 or ARPE-19 cells, in the presence or not of 2.5μM Ganciclovir. In MRC5 cells, infection progresses to reach more than 70% infected cells 10 days pi and this infection is partly controlled by Ganciclovir. ARPE-19 cells do not seem to be highly permissive and 2 to 3% of the cells only become infected, in the presence or not of Ganciclovir; d) same experiment as in c) but using viral DNA quantification as a read-out.

## DISCUSSION

In this study, we describe the first application of the ANCHOR DNA labeling technology to a virus resulting in ANCHOR-modified HCMVs which allowed follow up and quantification of HCMV infection and replication kinetics in real time and living cells, from few hours pi. until lysis of the infected cell. The biology of ANCHOR-HCMVs and parent BAC derived viruses appears similar with the replication rate of TB40-ANCHOR3 being even more robust (Table2, Fig.3). This increased replication rate may stem from variations in viral stock titration, but we may also have selected a virus especially fitting in MRC5 cells during the BAC construction which comprised cloning and low efficacy transfection steps. However, this TB40-ANCHOR3 virus seems to behave as expected and to be therefore an acceptable and representative HCMV with no significant compensatory deletion that have been described to sometimes affect HCMV-BAC overlong genomes (48). Moreover, TB40-ANCHOR3 viruses are tegumented, enveloped and contain viral DNA, possibly associated to OR-GFP proteins (Fig.4). It may seem surprising that a number of OR-FP sufficient to detect a fluorescent spot enters the capsid because the viral genome is generally supposed to occupy the entire inner volume of the capsid (49, 50, 51). This number has been estimated around n=50 (52) while the number of OR-GFP proteins bound to a single ANCH site and around was evaluated to n=+/-500 by FCS (23) on a chromosomal site integrated into the genome of a human cell. Using 1.15 nm (half the distance between concentric layers of packed DNA in ref. 51) and 0.85 10^5^ nm respectively as radius and length of a 250kb viral DNA, the calculated volume of the viral DNA (=3.53 10^5^ cubic nm) is less than the volume of a capsid with a internal radius of 48-50nm (49) (=4.63 10^5^-5.25 10^5^ cubic nm) and the remaining volume could easily accommodate up to 500 OR-FP proteins that would only occupy an estimated volume of 5.25 10^4^ cubic nm. Therefore, volume considerations do not preclude the presence of OR-GFP proteins inside the capsid even if the mechanism governing their introduction into the capsid remains unknown. Notwithstanding, the packaging mechanism which creates very high pressure inside the capsid and is able to adapt the compaction of the DNA to the length of the genome is not really understood (50). Of note is that OR-GFP can not enter any viral particle (results to be published elsewhere) indicating that each viral family has its own packaging specificities. ANCHOR-HCMV viruses are remarkably stable because >90% of particles conserve the ANCHOR phenotype through a 10^9^ amplification step. Similar results were obtained with TB40-GFP and non-fluorescent particles could thus simply reflect lack of expression or partial loss or mutation of any part of the ANCHOR insertion. Further experiments are needed to discriminate between these two possibilities. It was firmly established in yeast, drosophila and human cells that OR-FPs bind to their specific target sequences and form fluorescent spots that are easily visualized by fluorescence microscopy (21, 22). In this paper, we demonstrate that this is also true in ANCHOR-HCMV infected cells. While ChIP experiments have shown that OR-GFP proteins specifically bind to the ANCH sequence of the viral genome, spots are only observed when cells infected with an ANCHOR-HCMV virus are also expressing the corresponding OR-FP protein. The same OR-FP protein in presence of a non-ANCHOR virus genome does not form any spot. Therefore, the spots we observe in this study each represent a cluster of OR-FPs proteins specifically complexed to ANCH (and surrounding) sequences inserted in viral genomes, be they unique or in a concatemeric form, nude or encapsidated.

The first sign of infection of MRC5 cells by ANCHOR-HCMV is appearance of diffuse fluorescence in the entire cell about 4 to 5 hours pi., attributable to OR-GFP expression. This fluorescence increases gradually, mainly in the cytoplasm, until about 16h. pi. when one or few discrete spots, likely corresponding to the original incoming viral genomes, become visible in the nucleus. From this moment, the complete course of infection can be observed. These initial spots first multiply in small specific territories within the nucleus, called pre-replicative sites. As this first modest multiplication occurs rather early during infection, it is not clear whether the viral polymerase is already expressed and active at this stage or whether this amplification is performed by cellular polymerases. In HSV1 infected cells, inhibitors of viral DNA replication block the formation of RC but not of similar pre-replicative sites (53, 54). Interestingly, this also seems to hold true for HCMV as TB40-ANCHOR3 infected MRC5 cells only present pre-replicative structures but not mature RC when treated with Ganciclovir (Fig.S4). This initial pre-replicative stage is followed by a massive amplification step resulting in numerous (several hundreds) very intense spots which, after fusion of the different viral replication domains, result in a unique large nuclear compartment at about 72h pi. At this stage, only this compartment is precisely and uniformly stained with an anti-UL44 antibody and thus contains the polymerase associated processivity factor characteristic of the replication compartment (RC). Interestingly, ANCHOR technology allows quantifying fluorescence intensity of single spots and discriminating between background and viral genome associated fluorescence. Spot intensity is highly variable with the brightest spots being in the RC while those found outside the RC or in the cytoplasm seem homogeneous and weak. As shown in Fig.10, the diffuse GFP fluorescence background (in dark blue), grossly corresponding to the cytoplasm of the cell, may be assigned a value of 90 AU and is scattered with spots (in pale blue) corresponding to +/-120AU. The RC is also delimited by a line of spots of the same color. In the RC itself, colors corresponding to higher fluorescence values do not form a continuous gradient but vary by discrete steps, separated by +/-30AU. As each spot corresponds to one or more viral genome(s) and each viral genome contains a single ANCH target sequence which binds and recruits similar number of OR-FP protein, we assume that the stepwise variation of the fluorescence intensity correlates with the number of viral genomes present in each spot. Because spot intensities in the RC vary by steps of 30AU which is also the value between the background and the less intense spots, it seems logical to speculate that 30AU is the quantity of fluorescence generated by the OR-GFP proteins associated to a single viral genome. Therefore, spots of 120, 150, 180, 210 and 240 AU could correspond to 1, 2, 3, 4 and 5 viral genomes when correcting for the 30AU fluorescence background. In the RC, this interpretation makes sense to concatemers of varying number of viral genomes generated by a “rolling circle” replication mechanism and that will subsequently be cut by the terminase complex after encapsidation of a single genome unit (55). This “rolling circle” mechanism of lytic replication, similar to phage replication and widely accepted, has never been formally proven but very nicely fits with our data even if we cannot exclude other mechanisms involving θ structures for instance (1, 56). If this assumption is correct, low intensity RC, non-RC nuclear and cytoplasmic particles are probably capsids containing a single viral genome but at different stages of maturation while bright spots present in the RC and displaying between 150 and 240 FU are replicative structures harboring between 2 and 5 viral genomes. As already mentioned, the most intense spots are preferentially localized in the center of the RC and the fluorescence decreases towards the edge in concentric zones of discrete value (Fig.9c and d). A similar distribution of spot intensities has also been observed in ARPE19 cells (Supplementary Fig.S3) suggesting that the RC is highly organized with active replication occuring in the center of the RC and that this organization is a general feature of HCMV replication in infected cells. Then, the produced concatemeric structures could migrate from the center towards the periphery of the RC and would progressively be shortened as they encounter the preformed empty capsids of which the terminase complex cuts out unit long complete genomes which are encapsidated. Fig.9c suggests that this migration/encapsidation process arrives to completion at the nuclear membrane which is underlined by a brim of single viral genome spots. Finally, considering that most of the spots are in the RC and associated with 1 to 5 genomes, one can assume that such an infected cell contains at this precise moment 3000 to 5000 viral genomes. It is noteworthy that qPCR titration of viral genomes yield values ranging from eight to twenty thousand viral genomes per infected cell (Table 2).We consider these two sets of values based on spot counting and fluorescence intensity remarkably coherent especially as they correspond to instantaneous (fluorescence) and cumulative (qPCR) measurements.

Once synthesized, the viral genomes have to enter capsids which are assembled in the nucleus from the different capsid proteins imported from the cytoplasm. With their portal complex, capsids are able to cleave the genome concatemers resulting from the rolling circle mechanism and to internalize a single unit length genome through an ATP dependant mechanism (39). When analyzing the RC by correlation fluorescence/electron microscopy, unexpected results were obtained, suggesting that the classical capsid classification in A, B and C forms is possibly oversimplified. By analyzing numerous pictures (Fig.8 and results not shown), it appears that capsids that could be classified as C forms are rare and much less frequent than described (41). On the other hand, some capsids containing dense fragmented material are also clearly visible on Fig.8c. Of course, all these results could simply reflect the different conditions of infection and sample treatment, but could also indicate that more capsid forms exist which need to be understood. Whatever their significance, the three capsids on the left of figure 8d (marked with white arrows) are associated to three spots of equal intensity, but less intense than the spot on the right of the same picture. This association could of course be fortuitous but it is also tempting to speculate that these three spots represent capsids which have internalized one single viral genome and which are ready to leave the RC. The fourth capsid which appears empty in electron microscopy is in close contact with a stronger fluorescent signal and could thus be associated with a replicating, concatameric structure. However, the three capsids showing weak fluorescence resemble type B capsids, considered to be devoid of genetic material. This interpretation therefore remains highly speculative but, on the other hand, simple random association is also hard to imagine. As Fig.8c suggests there may exist other intermediate forms of capsids, it is possible that type B is a heterogeneous population of which a fraction could contain DNA.

Once capsids have loaded a viral genome, they leave the RC and the nucleus toward the cytoplasm. ANCHOR technology allows precise counts of the genomes present in different parts of the cell as illustrated in Fig.9. Due to their fluorescence values, we think that spots present in the nucleus outside the RC and in the cytoplasm correspond to single encapsidated genomes contrary to spots observed in the RC. If this interpretation is correct, there are significantly less viral genomes in the non-RC nucleoplasm and the cytoplasm than in the RC, possibly around 1% or less. Interestingly, when titers of virus stocks are plotted against the number of producing cells, obtained numbers are closer of what is found in the cytoplasm than in the RC, suggesting that most of the genomes synthesized in the RC could be lost. Therefore, the passage from the RC towards the cytoplasm could be a significant “bottleneck” in virus production even if the very low number of spots observed in the non-RC nucleoplasm could argue for the RC exit being the real limiting step. It is also possible that once encapsidated with the viral genome, fluorescence intensity of OR-FP is reduced and that only a minute fraction of the mature viruses remain visible. Reduced fluorescence of the encapsidated viral genome could be due to loss of OR-FP molecules during the encapsidation process. Cytoplasmic spots seen in Fig.9 are coherent with this hypothesis as they are clearly detected in Fig.9 a and b due to their fluorescence although they do not reach the threshold of 120 fluorescence AU. However, a cluster of 50-100 OR-FP proteins is already detectable and it is therefore logical to find in the cytoplasm spots with intensities ranging between background and 120AU. This may also explain why we were so far unable to visualize incoming viruses during the very first hours pi. although these could simply have been missed due to the extremely low probability to detect them in the adequate focal plane. A large proportion of the spots found in the cytoplasm are preferentially gathered in a faintly demarcated region close to the nucleus. As this region is stained by anti-pp28 and anti-gB antibodies (Fig.7), it very likely represents a viral Assembly Compartment where the capsids acquire their tegument and envelope and which overlaps the Endoplasmic Reticulum-Golgi-Intermediate Compartment (ERGIC) (35, 36, 37). In the rest of the cytoplasm, spots are fainter and their observation necessitates boosting the image acquisition conditions, leading to RC signal saturation. Finally, the way new viruses leave the cell remains so far unclear and possibly, particles could also be directly transferred from one cell to another neighboring one without any extracellular step. Alternatively, time lapse imaging of infected cells also revealed sudden “blebbing” and fragmentation of infected cells. This “blebbing” is often considered as indicative of apoptosis (38) but the mechanism of fragmentation ending cell infection is poorly understood. As the timing of this fragmentation is significantly delayed by the potent suppressor of apoptosis vMIA encoded by the UL37×1 viral gene, it is likely that longer infection time is advantageous for the virus and that fragmentation, on the contrary, interrupts the viral cycle and allows release of alarm signals to neighboring cells (57). Interestingly, this ultimate step occurred at very different times in different cells and generates numerous highly fluorescent cell fragments which thus possibly contain viral DNA. It will be of great interest to further explore this process to determine whether it may contribute to virus dissemination or whether it is the ultimate cellular defense bypassing the final maturation steps of the viruses and releasing non infectious although immunogenic material.

From a biotechnological point of view, ANCHOR-HCMV viruses are a remarkable tool to characterize antiviral compounds. As a proof of concept, we have measured the effect of Ganciclovir on the infection of various cell lines with ANCHOR3-HCMV. With very limited hands-on investment, we have been able to establish IC50 and IC90 of this drug on the HCMV infection of MRC5 human fibroblasts, simply recording fluorescence variation using an automated Arrayscan microscope (Fig.10). The results we obtained were highly reproducible and coherent with values already published (47). We performed the same experiment on the retina epithelial ARPE-19 cell line, with totally different results. These cells are indeed much less efficiently infected by the ANCHOR-HCMV TB40 virus, and the course of infection does not seem to be modified by Ganciclovir, suggesting that the drug is not metabolized in the same manner in MRC5 and ARPE-19 cells. For instance, phosphorylation of this drug by cellular kinases may be less efficient in ARPE-19 cells. Alternatively, higher drug concentrations may be needed as suggested by the loss of HCMV replication in these cells in the presence of 12.5µM Ganciclovir. Whichever the reasons underlying the different responses of different cell lines to Ganciclovir, these results indicate that ANCHOR-HCMV viruses will permit rapid and cost effective screening of large libraries of chemicals for the search of new anti-viral activities, including measurements of several parameters as toxicity of the compound, infection rate, virus DNA replication level and infection propagation without any fixation, extraction or reagent. Using ANCHOR-HCMV and automated high content microscopy, it will be easy to screen even large collections of chemical compounds. Furthermore, this technology can be used to label other DNA viruses for which we already have proofs of concept (results to be published elsewhere).

In addition to new insights into fundamental biology of numerous DNA viruses, ANCHOR technology is amenable to high throughput imaging, with high confidence over long time series of multiple cell lines, under different biological conditions in parallel. The technology is therefore particularly suited to rapid testing compound concentrations, stability and administration conditions for the design of new and/or combinatorial antiviral treatments. For all these reasons, the ANCHOR technology appears as a highly promising tool for fundamental research but also for numerous biotechnology applications.

## ACKNOWLEDGMENTS

We thank Eva Borst and Martin Messerle for the CMV TB40-GFP BAC construct and various plasmids. We greatly benefitted from the expertise of Sylvain Cantaloube at the TRI Imaging platform (IBCG) and of Jacques Rouquette at the ITAV imaging facility. We are also deeply indebted to Professor Henri Agut for critical reading of this manuscript. This work was supported by “Fondation ARC”, grant n° HFSPO RGP0044, and by a development project of “Toulouse Tech Transfer”.

